# Evidence for stabilizing selection at pleiotropic loci for human complex traits

**DOI:** 10.1101/126888

**Authors:** Emily S Wong, Steve Chenoweth, Mark Blows, Joseph E Powell

## Abstract

How genetic variation contributes to phenotypic variation is a central question in genetics. Association signals for a complex trait are found throughout the majority of the genome suggesting much of the genome is under some degree of genetic constraint. Here, we develop a intraspecific population genetics approach to define a measure of population structure for each single nucleotide polymorphism (SNP). Using this approach, we test for evidence of stabilizing selection at complex traits and pleiotropic loci arising from the evolutionary history of 47 complex traits and common diseases. Our approach allowed us to identify traits and regions under stabilizing selection towards both global and subpopulation optima. Strongest depletion of allelic diversity was found at disease loci, indicating stabilizing selection has acted on these phenotypes in all subpopulations. Pleiotropic loci predominantly displayed evidence of stabilizing selection, often contributed to multiple disease risks, and sometimes also affected non-disease traits such as height. Risk alleles at pleiotropic disease loci displayed a more consistent direction of effect than expected by chance suggesting that stabilizing selection acting on pleiotropic loci is amplified through multiple disease phenotypes.

## Introduction

There has been a long-standing interest in understanding how natural selection shapes genomic variation and facilitates adaptation in modern human populations (Sabeti et al. 2006; Vitti et al. 2013). The genetic architecture of human traits reflects the outcome of natural selection, demographic history and genetic drift. Advances in high-throughput genomics have revealed that many disease, anthropomorphic, and behavioral phenotypes are determined by a combination of alleles at a large number of independent loci of small effect size (Boyle et al. 2017; Yang et al. 2010). In line with this, based on the idea that adaptation is likely to be often polygenic, population genetics has focused to incorporated models of adaptive evolution from standing variation (Pritchard et al. 2010; Przeworski et al. 2005; Hernandez et al. 2011; Hermisson and Pennings 2005).

The study of polygenic adaptation has largely focused on the detection of responses to directional selection through soft selective sweeps (Messer and Petrov 2013; Field et al. 2016; Schrider and Kern 2017). However, to understand how standing variation in complex traits is maintained, a more intense research effort needs to be devoted to quantitating and understanding stabilizing selection. Stabilizing selection is difficult to study empirically but is central to a major class of theoretical models, the mutation-selection balance models, to explain quantitative genetic variation. Indeed, three decades of empirical research using phenotypic methods (Lande and Arnold 1983), has failed to reveal the abundant stabilizing selection that is predicted by theory (Kingsolver et al. 2001). Human genetic trait data from GWAS presents an unique opportunity to develop new approaches to gain a deeper understanding of the nature of stabilizing selection based on the underlying genetic architecture.

A key consideration to explaining genetic variation in polygenic traits lies in understanding and modeling pleiotropy (Johnson and Barton 2005). In humans, a number of methods have been recently developed to identify pleiotropic loci using GWAS datasets (Deng and Pan 2017; Pickrell et al. 2016; Wang et al. 2015; Han et al. 2016; Yang et al. 2017), and recent work has shown that pleiotropy is likely widespread across human complex traits (Pickrell et al. 2016; Visscher and Yang 2016). Given that the genetic variation in many, if not most, traits is likely determined by an accumulation of small effects across the genome, a key challenge lies in understanding how phenotypes independently evolve in the face of widespread pleiotropy (Lande 1980). Fisher’s geometric model of adaptation (Fisher 1930) predicts pleiotropic variants are less likely to be beneficial, and so for populations close to their phenotypic optimum, pleiotropic variants should experience stronger stabilizing selection (Orr 2000; Welch and Waxman 2003). Indeed, several trait values in human populations appear close to phenotypic optimum as stabilizing selection has been detected at the extreme tails of the phenotypic spectrum (Sanjak et al. 2018). However, evidence for the strength of stabilizing selection increasing with the extent of pleiotropy is limited. In one mutation accumulation experiment in flies, deleterious mutations that affect more than one gene expression trait are subject to stronger stabilizing selection (McGuigan et al. 2014).

Recent genome-wide association studies (GWAS) and large-scale genetic variant information provides an ideal resource for comparing the role of mutations in potential adaptive changes for complex phenotypes In the past decade, there has been rapid growth in methods to detect adaptive changes in humans at complex traits (Berg and Coop 2014; Galinsky et al. 2016; Field et al. 2016; Pickrell et al. 2009; Zeng et al. 2018; Sanjak et al. 2018).

To what extent pleiotropy may act under selective constrains at candidate loci for human complex traits has been little investigated. Here, we use a new intraspecific population genetics approach that allows us to specify a rigorous null model to assess both directional and stabilizing selection based on allele frequency change. We use this on the 1000 Genomes Phase 3 data to understand adaptive changes at complex trait loci using genome-wide association (GWA) summary statistics. We sought to address the following questions: (1) Does pleiotropic loci show evidence of directional selection? (2) What is the evidence for population-specific selective constrains at complex traits?

## Results

### A framework to test for selective constraints at genetic variants

For large SNP genotype data sets, principal components analysis can be used to detect population structure (Price et al. 2006). Genetic variants that are outliers in their contribution to the axes of genomic population structure are candidates for local adaptation. Under the assumption that most variants are evolving under a neutral model, we estimate population structure in the genomic relatedness matrix using random matrix theory to assign statistical significance to the principal axes of genetic variation. This forms the basis of a null model to identify SNPs whose allele frequencies are inconsistent with random drift, implying the action of natural selection. We test for evidence of natural selection acting on pleiotropic loci for complex anthropomorphic, behavioral and disease traits, under the expectation that trait-associated variants that are consistently under-represented or over-represented in axes of genomic population structure are candidates for local adaptation.

Specifically, we used matrix decomposition to devise a differentiation score (*d*) that identifies signatures of natural selection in extant human populations, and assessed the statistical significance of variants associated with complex traits under directional selection. Our method is most similar to that of (Duforet-Frebourg et al. 2016; Galinsky et al. 2016) and our d-score is similar to the calculation of the communality statistic of Duforet-Frebourg. However, our method differs in several key aspects. (1) Our method provides an unbiased, genome-wide measure with a null model for significance testing, lacking in many selection scans that examine outliers without a neutral model (Akey 2009). Only unlinked and common SNPs in Hardy-Weinberg equilibrium were used to estimate population structure. Including linked and rare SNPs, would violate our assumption of the population structure under neutrality and this estimation is the basis of the null model. We judge the significance of the departure of complex traits from neutrality by comparing the allele frequencies of SNPs from GWAS traits to the null distributions constructed from well-matched control SNPs. We note also, from a practical standpoint, most recent human genetic variant data, including the 1000 Genome Phase 3 cohort used here, are SNP dense; therefore the use of unlinked variants is the most computationally efficient strategy. (2) We use a statistical model to rigorously assess the number of axes of variations that contribute to population structure placing the metric on robust statistical grounds.

We used a model based on the Tracy-Widom distribution to define the number of eigenvalues that demarcates true population structure (Patterson et al. 2006; Bryc et al. 2013). A *d* score, measuring allelic differentiation, was computed for each single nucleotide polymorphism (SNP) (*n*=~12 million SNPs) (**Methods, FIG 1A-D, S1**). This is a measure of evolutionary change between populations, and corresponds well to the common measure of genetic differentiation – the *F_ST_* index (**FIG 1D**). We assumed the majority of genetic variation fits a neutral model (Miura et al. 2013), and that usually large or small values of *d* suggest the action of natural selection. High *d* indicates a change in allele frequency between orthogonal axes of relatedness variation, suggesting the action of positive selection or stabilizing selection within subpopulation/s to different optima. Alternatively, a low *d* value indicates the SNP does not contribute to population subdivision suggestive of stabilizing selection to a common optimum across all subpopulations. Compared with *F_ST_, d* has several advantages. *i)* Population differences are captured in a single framework accommodating multiple subpopulations, reflecting gradual evolutionary divergence and avoiding the pre-specification of population groups and accounting for admixed individuals *ii)* Population structure can be quickly estimated using genomic data from the whole genome *iii)* The measure can be calculated in one step for all commonly segregating SNPs, allowing quantitative assessment of evolutionary change at complex trait loci.

**Figure 1.**
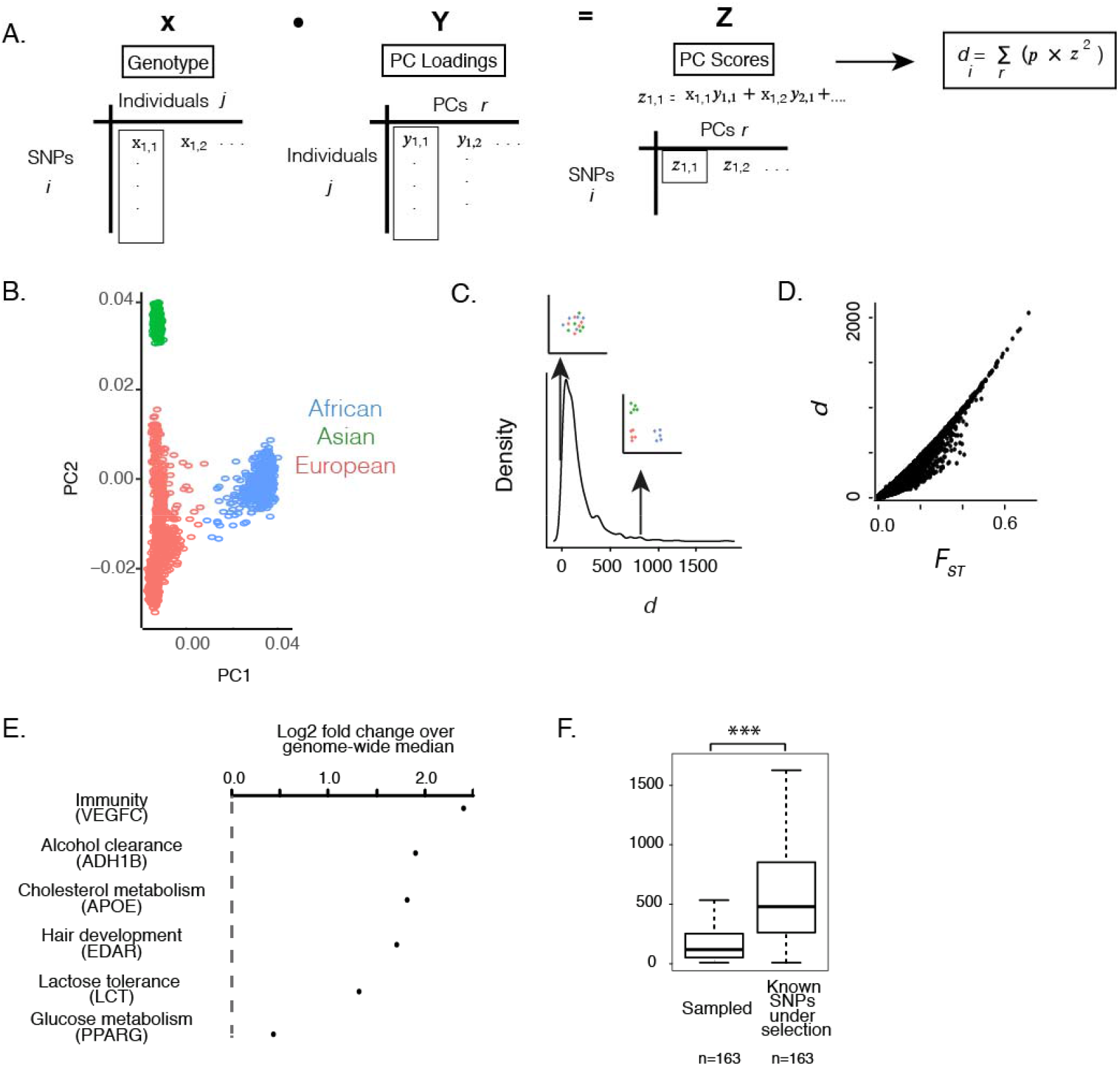
A measure of allelic differentiation (*d*) between human populations. (**A**) We used PCA to estimated allelic differentiation between populations within the 1000 Genomes Phase 3 dataset. For the top five PCs, the dot product of the standardized genotypes (**X**) and PC loadings (**Y**) and were used to calculate the PC score for each variant (*z*), which was squared and multiplied by *p_r_*, the proportion of variance explained by the rth PC. The number of eigenvectors used (the five largest eigenvalues) was determined using mathematical modeling to find the number of eigenvalues denoting non-random structure. (**B**) The first two PCs separate three major ethnicities comprising of 2504 individuals. (**C**) The resulting index, *d*, is a measure of genetic differentiation, where *d* increases with the degree of differentiation between populations. At high *d*, allelic differentiation at a SNP is high (as illustrated by the scatterplot where the colours represent grouping of individuals based on allelic difference). At low *d*, allelic differentiation at variants is low (population groups cannot be distinguished based on the information at that allele). (**D**) *d* is strongly linearly correlated to a common measure of population differentiation, *F_ST_*. (**E**) High levels of population differentiation can suggest positive selection driving changes in allele frequencies. Enrichment of *d* over genome-wide median is observed at well-known examples of loci that have undergone positive selection. (**F**) Increased *d* was observed at 120 loci previously reported to show high *F_ST_* between populations (Akey et al. 2002). The boxplot shows the distribution of *d* at randomly sampled SNPs (left) and at the previously published variants (right). The number of sampled variants were matched to the numbers presented from the published study.

To verify our approach, we replicated evidence for local selection at loci responsible for skin and hair pigmentation in genes *SLC24A5*, and *SLC45A2* responsible for light skin in Europeans, and at immune and metabolic genes (**FIG 1E-F**). Skin associated loci, including variants associated with gene expression, has been implicated in other tests for local adaptions selection (Fraser 2013; Field et al. 2016).

Given the availability of whole genome sequence data, we take advantage of the opportunity to examine regulatory regions and compared differences in *d* between functionally annotated regions and between loci associated to Mendelian traits and diseases. We quantified the overlap of 1000 Genome Phase 3 variants with a compendium of genomic locations of different functional annotations from published analyses (**Table S2**) Consistent with expectation, coding regions were the least genetically differentiated between populations indicative of strong purifying selection at these regions, while enhancer regions were often more genetically differentiated (**FIG S2**). The highest *d* scores, suggestive of increased differentiation between populations, were functionally enriched for genes and regulatory elements associated with skeletal structure, hair patterning, facial morphology, and glucose metabolism (**FIG S3**).

### Detecting natural selection at disease and anthropomorphic traits

To leverage the availability of GWA studies of complex traits and diseases, we compared GWA associated variants using summary statistics of 47 disease, anthropomorphic and behavioral phenotypes for differences in levels of *d* (**Methods**). Based on the assumption that the empirical distribution of SNP data from human populations is a good fit to a model of neutral mutation (Miura et al. 2013), we test whether *d* values of GWAS SNPs departed from the bulk of nucleotide polymorphisms in human populations that we assume to be well explained by neutral mutations.

To test the robustness of these results, we defined a distribution of null *d* values for each set of GWA variants on a trait-by-trait basis, and tested for significant departure of allelic differentiation from the null, using randomization and resampling to test for increased or reduced levels of genetic differentiation, controlling for type I errors. Our empirical null distribution was generated per trait based on resampling an equivalent number of SNPs to the number of independent/linkage disequilibrium (LD) filtered SNPs. Sampled SNPs were matched based on the joint distribution of the distance to transcriptional start site (TSS) and the minor allele frequency (MAF) to account for differences in the underlying evolutionary rates. The mean *d* from permutations was compared to the mean *d* of observed SNPs and was used to calculate an empirical *p*-value. (**Methods**).

Allelic differences suggesting the action of natural selection were most pronounced at disease loci. Depletion of genetic variation at disease-associated regions is most plausibly explained by the widespread selection against harmful mutations in a healthy cohort such as the 1000 Genomes (1000 Genomes Project Consortium et al. 2015). Regions associated with ulcerative colitis, rheumatoid arthritis, Alzheimer’s, type I diabetes (T1D), low-density lipoprotein levels and height showed reduced allelic differences suggestive of stabilizing selection (**FIG 2, S4**). In contrast, type 2 diabetes (T2D) and fasting insulin levels showed increased genetic differences between populations suggestive of selective differences between populations. Indeed, risk versus protective allele direction reveals consistency at disease traits associated with low *d*. The minor alleles at all genome-wide significant SNPs for T1D are more concordant between AFR vs EUR than expected by chance (binomial test p=1.1e-145, n=1183). On the other hand, for T2D where *d* values were high, there is no significant match between risk alleles in the two populations (binomial test p=0.83, n=89). The difference between the two traits is significant (Fisher’s Test p=1.1e-35).

**Figure 2.**
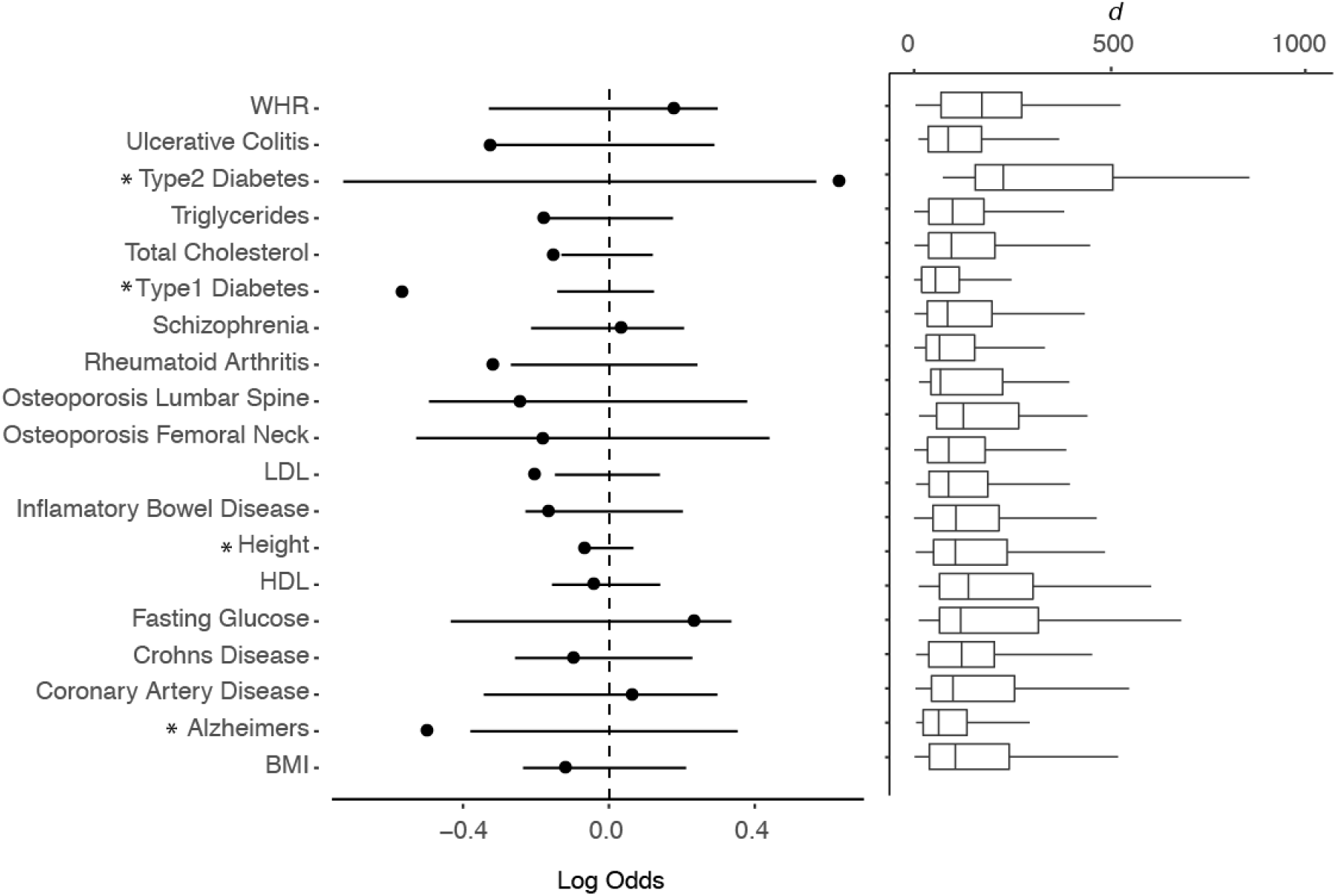
Genetic differences between populations (*d*) for complex trait loci. Higher and lower than expected levels of population differentiation suggests the action of natural selection driving changes in allele frequencies. Dot denotes the log odds calculated by dividing the median *d* values at genome-wide significant SNPs to the median of the matched null (averaged across 1000 permutations). Lower and upper quantiles are denoted by the start and end of horizontal lines. These values are medians of sampled *d* values from the specific background distribution for each trait. Boxplots describe observed *d* values for each trait. Only traits where the number of independent genome-wide significant SNPs > 10 are depicted here. Significance at two-tailed empirical *p*<0.05 is denoted by *. We performed the same analysis with a different GWA *p*-value cutoff, 1×10^−4^, (**SM**), which found significantly reduced *d* values at ulcerative colitis, triglyceride level, total cholersterol level and HDL associated loci (*p*<0.05).

Although height loci were significantly less differentiated between Africans and other subpopulations (Mann-Whitney U p<2.2e-16)(**FIG S5**), increased population differentiation for height loci was observed in the principal components demarcating Europeans and East Asian consistent with evidence for increased population differentiation at height loci within European subpopulations (Robinson et al. 2015), (Mann-Whitney U p<2.2e-16) and between Europeans (Mann-Whitney U p<2.2e-16)(PCs 2 and 3, respectively).

### Pleiotropic loci are predominately associated with disease and are under stabilizing selection

Theoretical and empirical studies have demonstrated that many complex traits are due to numerous loci of small effects (Rockman 2012; Pritchard et al. 2010; Pritchard and Di Rienzo 2010) implying many genetic loci are pleiotropic or shared between phenotypes (Gratten and Visscher 2016). Selection on genetic variants is a function of both on their effect on the focal trait and their pleiotropic effects on other traits (Simons et al. 2018). To examine selection at shared loci, we first used a Bayesian approach to determine pleiotropic loci based on cross-phenotype genetic associations of summary GWA statistics (Majumdar et al. 2017). Of the 7,344,447 SNPs where GWA effect size information was available for more than one trait, we identified 1,621 pleiotropic SNPs associated with 285 independent loci (log10 Bayes Factor>5) (**Methods**).

We tested for departure from background allele frequencies at pleiotropic alleles. Consistent with the action of stabilizing selection, we found significantly lower levels of *d* amongst pleiotropic loci compared to our null distributions of SNPs matched for MAF level and the distance to TSS (empirical *p*=5.6×10^−12^)(**FIG 3**). Most pleiotropic loci were linked to at least one disease, most commonly gastrointestinal disease, rheumatoid arthritis and type I diabetes. Among all GWAS traits studied, these traits displayed the lowest mean *d* values indicative of strong purifying selection against disease-causing genetic variants. As expected, many SNPs of large effect between phenotypes are located in the major histocompatibility complex (MHC)(Supplementary File X).

Next, we interrogated trait combinations where pleiotropic loci exhibit differences in selection pressure compare to their trait-specific backgrounds. For each pair of traits, we measured *d* at overlapping GWA variants removing SNPs determined to be in LD with one another (**Methods**). Significant departure of the *d* score from null was measured empirically by randomization and resampling of the equivalent number of tested SNPs on a per trait basis.

We found 11 unique trait pairs where shared SNPs displayed higher *d* and 15 trait combinations where *d* was reduced compared to trait background (empirical *p*<0.01)(**FIG 4**). A lower degree of allelic differentiation was observed at shared loci for years of formal education and conscientiousness, implying pleiotropic variants for these traits are selected towards a global optimum (empirical *p*<0.001)(**Methods**). Increased *d* indicative of population-specific differences in selection pressure was identified at shared loci for schizophrenia, education attainment and body fat associated traits (empirical *p*≤0.01) (**FIG 4**). We caution that a high *d* could be also influenced by subpopulation differences in causal loci.

**Figure 3.**
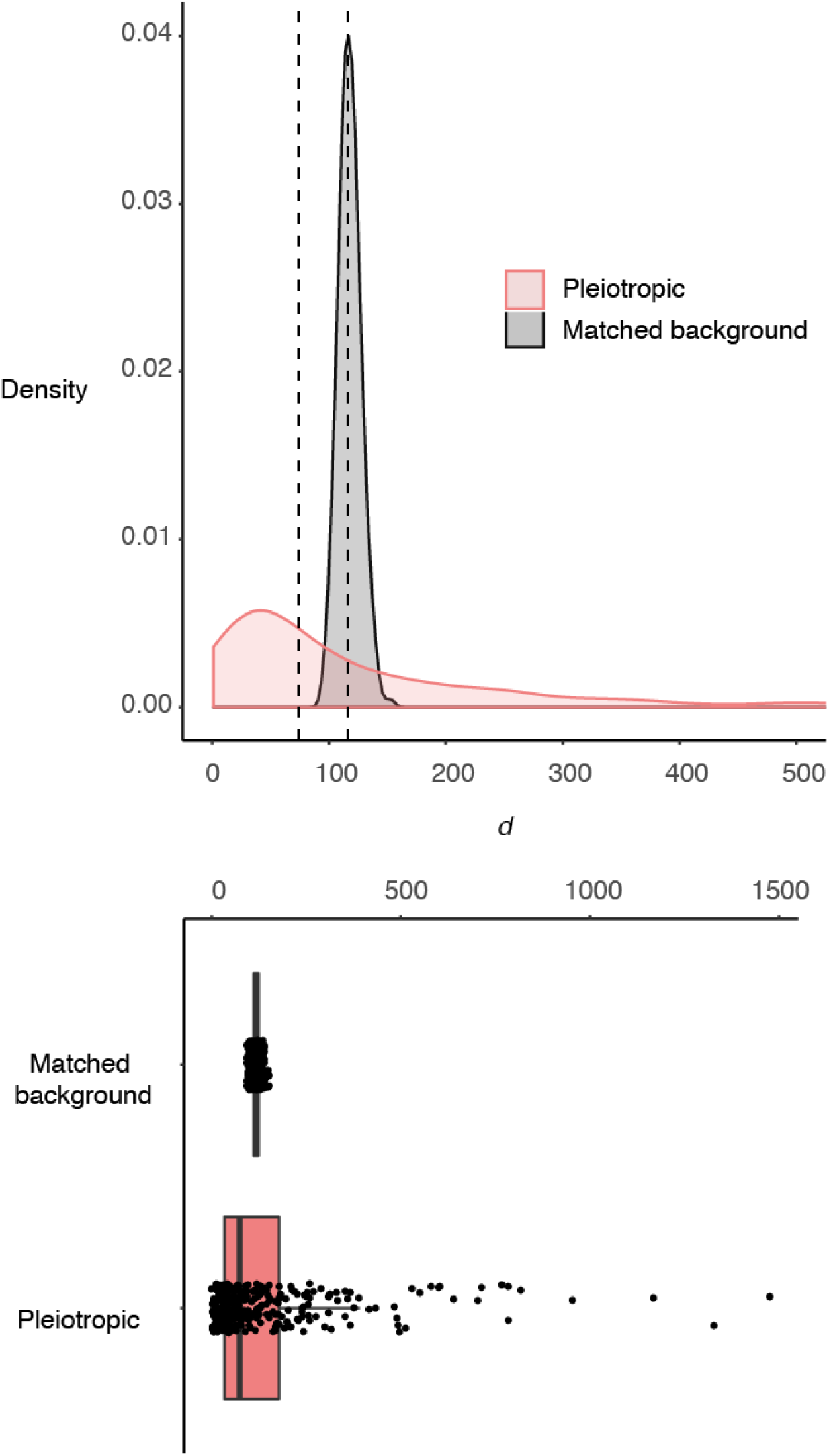
Evidence of purifying selection at pleiotropic loci. A reduction of *d* over background suggests the action of greater stabilizing selection at pleiotropic loci. Pleiotropic regions comprised of 285 unlinked loci (**Methods**). Permutations were used to assess the significance of the mean *d* value by sampling an equivalent number of SNPs across the genome matched for MAF and distance to TSS. This procedure was repeated 1000 times to calculated an empirical one-tailed *p*-value. Dashed lines denote the overall median values for pleiotropic and background SNPs matched for MAF and distance to TSS.

**Figure 4.**
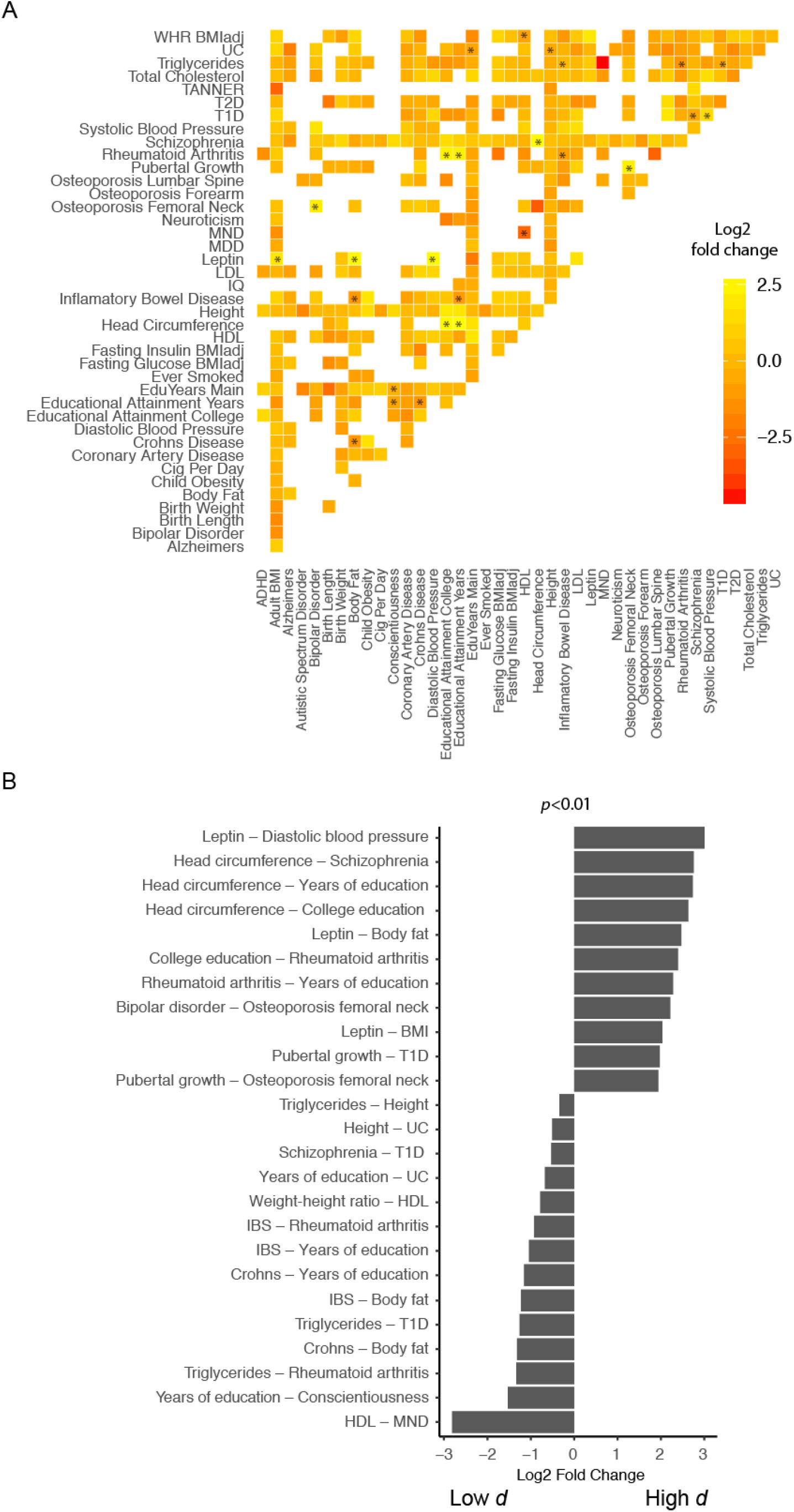
Pairwise trait comparisons reveal increased genetic constraint at shared loci for trait combinations. An increased or a reduced level of population differentiation suggests the action of selection at pleiotropic loci. (**A**) We first obtained shared variants between traits by taking the mean *d* of overlapping variants. Significance (two-tailed) was then calculated by comparing the mean *d* value at overlapping SNPs by sampling from trait-specific backgrounds for 1,000 permutations. The heatmap depicts change in *d*, log2 (mean expected *d* / mean observed *d*), from trait-specific background at pleiotropic loci. * denotes where shared SNPs for the two traits show an increased or decreased *d*, indicative of common regions that have undergone selection. Due to the polygenic nature of complex phenotypes, we quantitated allelic differentiation for all SNPs with GWA *p*-values below 1×10^−4^. (**B**) Barplots depict pairwise comparisons where shared SNPs show a departure from background values with *p*<0.01.

A valuable aspect of GWA summary statistic is the directionality of the trait effect. Pleiotropy can cause covariation among traits that can promote or constrain evolutionary change (Walsh and Blows 2009). The relationship between pleiotropy and genetic constraint is, however, not straightforward. When trait combinations are under selection the direction of evolutionary response in single traits can be unpredictable owing to nonlinearity of selection and complex genetic relationships between traits ^37^. To gain insights into the directional contribution of GWAS traits at pleiotropic loci, we considered how alleles shared between traits contribute to the common locus when traits are evolving under selection. We classified shared loci by genomic proximity, whereby GWA variants for all traits (*p*-value cut-off of 1×10^−4^) were clustered.

Applying this approach to examine the consistency of allelic direction at pleiotropic SNPs shared between traits revealed significant concordance of risk alleles (binomial test *p*=9.0×10^−11^, n=204). Therefore, stabilizing selection acting on a pleiotropic locus through one disease phenotype will tend to be reinforced by stabilizing selection acting through other disease phenotypes as a consequence of the positive genetic correlation generated by these loci.

Non-disease phenotypes may also share pleiotropic loci with disease phenotypes, and reinforce the action of stabilizing selection acting on disease phenotypes. This effect is exemplified at loci shared by height, a highly polygenic trait evolving under stabilizing selection with predicted non-null effects across the majority of common SNPs in the human genome (Boyle et al. 2017; Field et al. 2016), and disease traits. At genomic regions where height and a disease trait collocate, a height increasing allele is as likely to co-occur with a disease-increasing allele compared to non-risk allele (n=243).

We further observed pleiotropic variants were commonly associated among genetically uncorrelated diseases. For example, we identified regions affecting both inflammatory bowel disease (IBD) and rheumatoid arthritis (e.g. rs1265098); osteoporosis and IBD (e.g. rs12742784); schizophrenia and coronary artery disease (e.g. rs284861). Notably, these diseases commonly cooccur. Arthritis is found in as many as 15-20% of Crohn’s disease patients (Orchard 2012), osteoporosis is a common affliction of individuals with IBD, and schizophrenia patients have an elevated risk for coronary artery disease (Hennekens et al. 2005). Thus, shared genetic etiology characterized by pleiotropic regions may play a central role in co-occurring diseases.

#### Disparities in population differentiation between related complex diseases

Next, we examined genomic patterns of population differentiation between related traits. To quantitatively compare the degree of allelic differentiation between GWAS traits, we modeled the distribution of *d* values at genome-wide significant SNP for each trait using a gamma distribution and compared between pairs of traits using statistical models and a likelihood-ratio test (LRT) statistic (**Methods**).

In several instances, genetically correlated traits were significantly different in their distribution of *d* at genome-wide significant SNPs. For example, childhood obesity and adult BMI are positively genetically correlated (*r_g_*=0.7) yet show significantly different distributions of *d* at significant SNPs (LRT *p*=1.3e-6) (Bulik-Sullivan et al. 2015a). We also observed a similar pattern between HDL and triglyceride levels (*r_g_*=-0.6) (LRT *p*=1.2e-4) (Bulik-Sullivan et al. 2015a). Ulcerative colitis and Crohn’s disease were also significantly correlated (*r_g_*=0.54), but population structure appear reduced at ulcerative colitis loci (LRT *p*=3.7×10^−13^) (**FIG S6**).

We further examined the degree of genetic differentiation amongst populations for related traits type I (T1D) versus type II diabetes (T2D) and between ulcerative colitis and inflammatory bowel disease (IBD).

T1D loci compared to T2D loci showed remarkably different signatures of genetic differentiation (**FIG S7A**). T1D associated loci displayed much reduced levels of genetic differentiation across populations compared to T2D loci (**FIG S7B**, Mann Whitney U *p*=8.4×10^−5^). Decreased population differentiation is suggestive of extensive purifying selection while signatures of increased genetic differentiation at genetic loci indicate differences in allele frequencies across different populations. T2D loci with significantly increased population differentiation were proximal to four genes that have been robustly implicated in the disease. Earlier studies using HapMap variants have reported high genetic differentiation between populations at *TCF7L2* (Klimentidis et al. 2011; Pickrell et al. 2009), and we newly identified enrichment of genetic differentiation at variants associated with *PPARG, WFS1*, and *IGF2BP2*.

We further compared LD scores at loci associated with both diseases. LD scores provide a summary of linkage disequilibrium in a local region (Bulik-Sullivan et al. 2015b). Measures of LD, such as long-range haplotypes in a population, are commonly used to characterize the increased levels of LD expected in a region undergoing positive selection (Sabeti et al. 2002). We observed that LD scores were also more highly elevated for T1D versus T2D indicating stronger ongoing selection against deleterious T1D alleles (**FIG S7C**, Mann-Whitney U *p*=9.5×10^−6^).

Increased allelic differentiation at T2D loci in human subpopulations is likely due to a combination of genetic drift and positive selection where increased genetic differentiation at some T2D loci could indicate soft selective sweep. However, it is unknown whether these variants were directly selected, or indirectly so through pleiotropy and hitchhiking, where an allele is linked to the sweep of a beneficial allele. The *d* score does not inherently distinguish between the direction of the SNP (i.e. risk versus protective alleles) as it is taking the square of the PCA scores for each axis of variation, so differences in directions between SNPs are not visible in the score, only the magnitude of differences are. However, findings from tests for concordance between direction of risk alleles between populations found no significant correlation for T2D loci taken together suggesting that individual variants may have been indirectly selected for other advantages (Berg and Coop 2014). This result is consistent with evidence of adaptation of gene expression levels linked to cold climate tolerance at genes that are also involved in diabetes-related pathways (Fraser 2013).

Taken together, the results reveal surprisingly distinct patterns of population structure for loci of related diseases. These large differences in genetic architecture are suggests differences in the degree of selective pressure between related diseases.

## Discussion

These results demonstrate evidence for pleiotropic loci acting under stabilizing selection in human populations. Our analyses were performed using unlinked SNPs at complex traits, which potentially introduces a bias to the results. As increased LD is also a feature of regions under adaptive pressure, this can make the results more conservative. However, LD and allelic differentiation can elucidate different mechanisms of selection. Selection for increased LD can be result of certain genomic arrangements (i.e. conserved synteny) that promotes molecular efficiency within a genomic neighborhood. On the other hand, allelic differentiation, as investigated here, can be evidence of selection at point mutations that disrupt protein-binding sites or at transcription factor binding motifs and do not necessarily imply increased LD.

We have replicated our analyses across GWA studies using an independent cohort and obtained highly concordant results (**Method, FIG S8**). By examining empirical evidence, we found pleiotropic loci underpinning the genetic basis of traits are often under differing degrees of selective pressure contingent on the nature of the traits. For example, negative selection was detected for pleiotropic loci between years of education and conscientiousness yet shared loci for head circumference and years of education as well as rheumatoid arthritis and years of education showed increased population differences. The results suggest 1) within the same trait different molecular pathways are subject to different selection pressure 2) hitchhiking of genetic effects through pleiotropy is common 3) educational attainment reflects influences from different biological pathways, including immune-related pathways. By varying the weighting of the eigenvalues in *d*, it is possible to obtain further insights into how adaptations vary between human subpopulations. It is also possible restrict the analysis to focus on SNPs that are population-specific, enriching for more recent mutations.

Stabilizing selection is often amplified at pleiotropic variants through disease phenotypes, but consistent with reported findings of an irregular relationship of effect sizes at pleiotropic variants (Pickrell et al. 2016), we observed that the direction of effect is also often unpredictable.

The results also highlights that genetically correlated traits are not necessarily subject to the same selective constraints at genome-wide significant loci and is consistent with the fact that a high genetic correlation does not imply that the two traits cannot evolve independently to some degree. Indeed, artificial selection experiments indicate that bivariate correlations do not adequately capture the complexity of selection and even highly genetically correlated traits are able to independently evolve to some extent (Hine et al. 2014; Beldade et al. 2002).

Further work is required to understand the complex relationship between pleiotropy and selection. A linear model fitted between *d* and the likelihood of pleiotropy of variant only displays a very weak positive relationship (**FIG S9**). The underlying relationship between pleiotropy and allelic differentiation is challenging to model due to the many variables that influence it. For example, as noted in **FIG S2**, the extent of *d* varies with the biochemical function of the region under consideration. Protein coding regions function in a different way and evolve in a different manner to regulatory regions such as enhancers as reflected by lower *d* values. This mans that for pleiotropy loci located at enhancers negative selection (inferred by decrease in *d*) tends to be weaker than at protein-coding regions. While it may be useful to further partition pleiotropic SNPs into subcategories, the power for these comparisons is low due to reduced SNP numbers. In addition, we have defined pleiotropy loci based on the GWAS studies that are available; these traits are limited and are also not uniform in their degree of similarity. This means that pleiotropic loci should ultimately be compared taking into account the relatedness of traits under consideration, and we should also bear in mind unmeasured phenotypes when examining a variant currently labeled as not pleiotropic.

There are a few other caveats to interpreting the biological implications of this study. It is important to note that most GWAS have been performed in individuals of European ancestry. This means that although GWA SNPs discovered in one genetic background can often describe the same phenotype in another population of a different genetic background, loci identified for a GWA phenotype in one population, for example, Europeans, specifically describes differences among individuals in that population and the same loci in other populations may not reflect the same trait, evolutionary forces or demographic events. For instance, educational attainment (Okbay et al. 2016) and height (Wood et al. 2014) GWA studies were both based on mapping individuals of European descent and the interpretation of their associated genetic loci in other populations should be interpreted with caution. However, various studies performed on different populations have shown that associated variants tend to replicate across populations (de Candia et al. 2013).

### Accounting for population stratification

Our method, like others based on comparisons of genetic differentiation (Berg and Coop 2014), would be prone to the presence of population structure in the GWAS panel, as the presence of population stratification that is not adequately corrected can confound GWA results by producing false positives with high genetic divergence. This is a well-known problem, and most recent GWA studies use statistical, or experimental design strategies to correct for population stratification. Nevertheless, to access the potential impact of population stratification on our results we simulated a polygenic trait using a non-ascertained cohort (GERA) and show that standard PCA correction for population stratification in a GWA study is sufficient control for this bias in our study (*45*, **Methods, FIG S10, Table S3**). Notably, we also obtained highly concordant results between two height GWA studies where in one study (Robinson et al. 2015) effect size estimates were inferred from sib-pairs and is thus robust against stratification (data not shown).

Despite a limited selection of GWA studies, we recovered pleiotropic loci under selection for the vast majority of disease traits suggesting the number of truly independent traits under strong stabilizing selection is small. Taken together, these findings underscore how the evolution of a complex trait is entwined with the dynamic interplay of selective forces across a large number of genetic loci.

## Methods

### Study cohorts, data preparation and preprocessing

The primary analyses are based on 1000 Genomes Phase 3 data comprising of 2,505 individuals (1000 Genomes Project Consortium et al. 2015). Prior to estimating global population structure, we used PLINK 1.9 (Purcell and Chang; Chang et al. 2015) to remove SNPs with a minor allele frequency (MAF) less than 1%. SNPs that were not in Hardy-Weinberg Equilibrium were also removed (*p*<1×10^−6^) and the data was iteratively LD-pruned with an *r^2^* cutoff of 0.2. SNPs located on sex chromosomes were also removed. To test the robustness of our approach and to assess potential effects of population stratification on our results, we also ran similar analyses, with the same preprocessing steps, using published and simulated GWA data using the Genetic Epidemiology Research on Aging (GERA) cohort, a subsample of the longitudinal cohort of the Kaiser Permanente Research Program in the Northern California region, which contained four self-reported ethnic groups (European, East-Asian, Latin Americans and Africans). Details of the GWAS studies used in our comparisons are found in **Table S1**)

### Quantitation of selection through matrix decomposition

We used principal component analysis (PCA) to identify population structure amongst a cohort of individuals using genotype data. We orthogonally transformed genotype data from the 1000 Genomes Phase 3 cohort (n=2,505) (1000 Genomes Project Consortium et al. 2015), and determined the principal components (PCs) that were informative of population structure using the Tracy-Widom distribution (Patterson et al. 2006; Bryc et al. 2013).

If a centered genotype matrix **X** (where means are subtracted from each column) is of size *i* × *j*, where *i* is the the number of SNPs and *j* is number of individuals, then the *j* × *j* sample covariance matrix 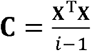 is the familiar genomic relatedness matrix commonly used in GWAS. This is a symmetric matrix and can be diagonalized:

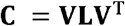

where **V** is a matrix of eigenvectors as columns, and **L** is a diagonal matrix with eigenvalues in decreasing order along the diagonal.

For numeric stability, PCA can be performed using singular value decomposition (SVD) of the genotype matrix **X**.

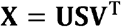

where **S** is the diagonal matrix of singular values *s_i_* Hence,

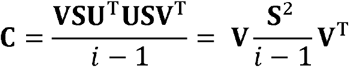

it is evident from this that **V** are the eigenvectors and the PC scores, **XV**, are equal to

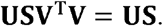

We used FastPCA (https://github.com/ajverster/FastPCA) to perform PCA using this approach. The method uses an approximation procedure of performing SVD by first randomly sampling the data into subspace based on the method of Halko *et al*. (Halko et al. 2011). The dataset was first preprocessed to remove variants that may bias the estimation of population structure. We used PLINK 1.9 (Purcell and Chang; Chang et al. 2015) to remove SNPs with a minor allele frequency (MAF) less than 1%. SNPs that were not in Hardy-Weinberg Equilibrium were also removed (*p*<1×10^−6^) and the data was iteratively LD-pruned with an *r^2^* cutoff of 0.2 (Galinsky et al. 2016). SNPs located on the sex chromosomes were also removed. The data was then mean centered and standardized to unit variance prior to PCA and PCA was run on 818,280 SNPs across individuals.

To identify the number of eigenvalues consistent with population structure, we used the eigenvalues of the uncentered and unstandardized covariance matrix, 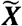, to infer the number of subpopulations in the data based on the method of (Bryc et al. 2013). The method analyzes the genotype matrix as a random perturbation of a finite-rank matrix. The number of significant eigenvectors, *r*, are estimated as the number of eigenvalues larger than the threshold of *t*′:

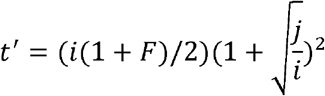

where *i* is the number of SNPs and *j* is the number of individuals. If *F_r,i_* is an inbreeding parameter for SNP *i* in subpopulation *r* that ranges from 0 to 1, then:

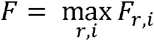

where *F_r,i_* = *F_r_* has the same value for all markers *i*, and *F_r_* is the average inbreeding coefficient of the *r*th subpopulation.

Given *F*=0, we find *r* = 6. This implies at least 6 subpopulations are present in the uncentered dataset. Centered data have *r*−1 large eigenvalues (Bryc et al. 2013). Hence, the first five eigenvalues from our centered matrix, ***X***, explaining 47% of total variance, are informative of population structure.

The matrix decomposition step above allowed us to estimate population structure among the *j* individuals. We now define a metric to reflect the extent of genetic differentiation between populations. To do this, we calculated **Z**, the *i* × *r* matrix of loadings, z_*i,r*_ for the *i*th SNP of the *r*th PC using:

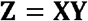

where **X** was the standardised genotype matrix, and **Y** was the *j* × *r* matrix of *y_j,r_* PC loadings for individual *j* for the *r*th PC. To obtain a measure of populations structure for all variants, including those removed due to linkage disequilibrium and those not in Hardy-Weinberg equilibrium, SNPs which were filtered out previously for the accurate estimation of population structure were reintroduced for this calculation.

The z_*i,r*_ provide information on how each SNP contributes to each of the subpopulations.. Finally, we summed the squared z_*i,r*_ values for each of the subpopulations into one score, *d_i_*, weighting each z_*i,r*_ by the amount of variance proportionally explained by the respective PC:

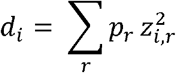

where *p_r_* is the proportion of total variance explained by the *r*th PC. A high value of *d_i_* indicates that the SNP, *i*, has changed frequency between the different orthogonal axes of relatedness variation (PCs), while a low value indicates the SNP does not contribute to population subdivision.

To examine selection within European populations, we used the same strategy to define a *d* score for individuals of European descent from the 1000 Genomes Project (number of individuals = 503; number of LD removed SNPs = 360,054). Based on the Bryc et al. 2013 test described above, we found only the first component demarcates population structure. Thus, we used the first PC to calculate a *d* score for each SNP.

### Significance testing of selection at GWA trait

We performed analyses for genome-wide significant SNPs but also extended our analysis to consider all GWA variants below a *p*-value <1× 10^−4^, as the majority of polygenic trait heritability is due to SNPs that do not reach genome-wide significance (Yang et al. 2010). At each functional annotation region for each trait, we LD pruned SNPs (*r^2^* cutoff=0.2) and calculated the mean *d* score for variants in linkage equilibrium. To control type I error, we used permutation strategies to assess the extent of increased or reduced levels of genetic differentiation. We performed 1000 permutations to estimate the significance of our results by resampling SNPs from the same functional category.

We calculated an empirical *p*-value based on generating a null distribution for each variant by subsampling based on matched MAF and distance to TSS. We first binned all 1000 Genome variants into 10 equally sized bins for MAF and 20 equally sized bins for distance to TSS. By sampling from matched bin sizes, we generated the background distribution for each variant and used this to calculate an empirical *p*-value. For each trait, we performed 1000 permutations to generate an empirical null by resampling matched number of SNPs to obtain 1000 values of median *d* expected and this was compared to the observed median *d* value.

A significant increase of *d* indicates selection is detected in at least one subpopulation. It is important to note that if the trait was studied only using individuals of one genetic background, it remains possible that an associated variant may not necessarily be linked with the trait in other subpopulations. Notwithstanding this possibility we note that concordance between GWA SNPs ascertained in different genetic backgrounds have commonly been observed (de Candia et al. 2013).

### Identification of pleiotropic loci and test for selective constraint

We used the R package ‘CPBayes’ (Majumdar et al. 2017) to identify genetic loci associated with multiple phenotypes using the GWAS summary data used above. Pleiotropic loci were determined based on a log10 Bayes factor cutoff 5. Both Crohn’s disease and ulcerative colitis are types of IBD, hence we have excluded instances where the most important phenotypes that underlie a pleiotropic signal involved only IBD and Crohn’s disease or only IBD and ulcerative colitis as we cannot be certain that these are true cases of pleiotropy. SNPs in LD with each other were removed using PLINK with similar criteria to above to identify independent pleiotropic loci. For each pleiotropic region (where there were two or more traits associated), we tested for departures of the averaged *d* against a background of resampled variants matched for distance to TSS and MAF. Significance (two-tailed) was calculated by comparing the mean *d* value for each pleiotropic loci by sampling from its null background for 1,000 permutations.

### Randomizations to assess the significance of genetic differentiation at shared loci between pairs of traits

In a pairwise manner, we assessed whether shared loci between traits possessed a *d* score that was significantly different from the trait-specific background from each trait. For each pair of traits, we overlapped GWA variants (*p*-value cut-off=1×10^−4^) removing SNPs determined to be in LD with one another based on the parameters described above. For each trait pair, we tested for departures of the mean *d* against an empirical distribution of means of sampled SNPs. The same number of SNPs, matched for distance to TSS and MAF, were sampled from the total pool of GWAS SNPs specific to the two traits under comparison. Significance (two-tailed) was calculated by comparing the mean *d* value for each pair of traits by sampling from its null background for 1,000 permutations.

### Framework to compare differences in *d* between GWAS traits

To compare trends in the distribution of *d* between GWAS traits we model the distribution of *d* using a gamma model,

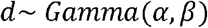

where *α* is the scale parameter and *β* is the rate parameter and the probability density function is defined as:

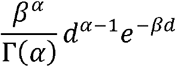

We performed all possible pairwise comparisons for each trait. To compare between traits we test the following models:

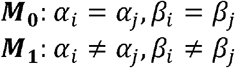

where *i* and *j* are the traits under comparison. ***M*_0_** denotes our null model where the density distribution is assumed to be the same between the traits. ***M*_1_** allows both parameters to vary between the traits. Maximum likelihood estimation was used to obtain *α* and *β* under ***M*_0_** and ***M*_1_**. We used a likelihood ratio test based on the chi-square distribution with two degrees of freedom to compare between ***M*_0_ and *M*_1_** P-values were adjusted for multiple testing using FDR correction and we considered q-values below 0.1 as denoting a significant difference in allelic distribution between traits. To compare between models, we also calculated the Bayesian information criteria (BIC).

### Assessing the impact of population stratification at GWA loci

Population stratification in GWA samples poses a concern if there is a match in the discovery GWA study between the direction of phenotypic differentiation among populations and genetic stratification in allele frequency in the 1000 Genomes dataset. Although this confounding effect is typically controlled for in discovery GWA studies, there is no guarantee that all stratification has been removed (Robinson et al. 2015). In such cases, inadequate control of stratification could lead to GWA variants with high levels of allelic differentiation between populations.

We performed simulations to assess the potential impact of population stratification in the discovery GWA study on our overall results. First, we simulated a polygenic trait using genotype data and use standard protocols for correcting population stratification to examine whether this is sufficient to remove false positives in our results. Next, given that some studies comprised only of European cohorts, we tested whether stratification in the discovery GWA study could systematically bias our results when stratified variants in the European population were projected to the 1000 Genomes population.

#### Simulation of a polygenic trait using an independent dataset with and without correction of population stratification

First, we randomly sampled 500 individuals from each of the four self-reported ethnic groups (European, East-Asian, Latin Americans and Africans) in the GERA (Genetic Epidemiology Research on Aging) cohort. The GERA cohort is a subsample of the longitudinal cohort of the Kaiser Permanente Research Program in the Northern California region. High-density genotyping was done with custom arrays for each of the four major ethnic groups. Next, we randomly assigned 5,000 independent loci across the genome to be causal variants. Their effects were sampled from a normal distribution with a mean of 0 and a variance of 1. The heritability of the trait (*h*^2^) was set at 50%. A phenotype was created as *y* = Σ *x_k_b* + *e*, where *x_k_* is the genotype copy at locus *k* (0,1,2), *b* the effect size 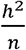, where *n* = 5,000 and *e* is the residual variance where *e* = *N*(0, 1 − *h*^2^).

We used PLINK 1.9 (Purcell and Chang; Chang et al. 2015) to estimate SNP effects using ordinary least-squares regression without any control for population stratification. We then repeated the GWAS estimation and controlled for stratification using the first three principal components as covariates in the regression. As expected, we found that correction for stratification reduced *d* values across the genome (**FIG S10**).

If the predictors are created from SNPs that were unbiased by population stratification, we expect no significant difference in overall measure of phenotype differentiation versus null expectation. This was observed to be the case (**Table S3**).

### Replication in an independent cohort

We replicated our study on the GERA (Genetic Epidemiology Research on Aging) cohort. We sampled 500 individuals from each of the four ethnic groups: European, East-Asian, Latin Americans and Africans. Analyses were repeated as described above for the 1000 Genomes dataset. Variants not genotyped amongst all selected individuals were filtered out. 6,690,254 variants remained after filtering (MAF>0.01). Following iterative pruning, 366,178 variants were used in PCA to estimate population structure.

We compared the log(observed mean *d*/expected mean *d*) for variants classed by GWA phenotype and annotation category between GERA and 1000 Genomes and observed close correspondence between results r=0.63 (Pearson’s correlation *p*-value<2.2×10^−16^) (**Fig S7**).

---

**Supplementary File 1: Supplementary materials including text, figures and tables**

**Supplementary File 2: *d* scores of variants in the 1000 Genome Phase 3 panel**

**Supplementary File 3: Pleiotropic loci, significance values, their associated traits, *d* score for each variant and genomic location (hg19)**

## Data accession

The GERA dataset was obtained from the database of Genotypes and Phenotypes (dbGaP) through accession phs000674.v1.p1.

## Contributions

ESW, SC, MB and JEP designed experiments. ESW performed analyses. JEP supplied data. All authors edited the manuscript.

## Acknowledgements

We would like to thank Matthew Robinson for his advice and input on use of the differentiation score. We would also like to thank Bruce Walsh, Paul Flicek, and Greg Gibson for valuable discussion and feedback on the manuscript. ESW is supported by an Australian Research Council Discovery Early Career Award (DE160100755). JEP is supported by National Health and Medical Research Council Career Development Fellowship (1107599) and grant (1083405). The authors have no completing interests.

## References

1000 Genomes Project Consortium, Auton A, Brooks LD, Durbin RM, Garrison EP, Kang HM, Korbel JO, Marchini JL, McCarthy S, McVean GA, et al. 2015. A global reference for human genetic variation. Nature 526: 68–74.

Akey JM. 2009. Constructing genomic maps of positive selection in humans: Where do we go from here? Genome Res 19: 711–722.

Akey JM, Zhang G, Zhang K, Jin L, Shriver MD. 2002. Interrogating a High-Density SNP Map for Signatures of Natural Selection. Genome Res 12: 1805–1814.

Beldade P, Koops K, Brakefield PM. 2002. Developmental constraints versus flexibility in morphological evolution. Nature 416: 844–847.

Berg JJ, Coop G. 2014. A Population Genetic Signal of Polygenic Adaptation ed. M. W. Feldman. PLoS Genet 10: e1004412.

Boyle EA, Li YI, Pritchard JK. 2017. An Expanded View of Complex Traits: From Polygenic to Omnigenic. Cell 169: 1177–1186.

Bryc K, Bryc W, Silverstein JW. 2013. Separation of the largest eigenvalues in eigenanalysis of genotype data from discrete subpopulations. Theor Popul Biol 89: 34–43.

Bulik-Sullivan B, Finucane HK, Anttila V, Gusev A, Day FR, Loh P-R, ReproGen Consortium, Psychiatric Genomics Consortium, Genetic Consortium for Anorexia Nervosa of the Wellcome Trust Case Control Consortium 3, Duncan L, et al. 2015a. An atlas of genetic correlations across human diseases and traits. Nat Genet 47: 1236–1241.

Bulik-Sullivan BK, Loh P-R, Finucane HK, Ripke S, Yang J, Schizophrenia Working Group of the Psychiatric Genomics Consortium, Patterson N, Daly MJ, Price AL, Neale BM. 2015b. LD Score regression distinguishes confounding from polygenicity in genome-wide association studies. Nat Genet 47: 291–295.

Chang CC, Chow CC, Tellier LC, Vattikuti S, Purcell SM, Lee JJ. 2015. Second-generation PLINK: rising to the challenge of larger and richer datasets. GigaScience 4. http://gigascience.biomedcentral.com/articles/10.1186/s13742-015-0047-8 (Accessed January 15, 2017).

de Candia TR, Lee SH, Yang J, Browning BL, Gejman PV, Levinson DF, Mowry BJ, Hewitt JK, Goddard ME, O’Donovan MC, et al. 2013. Additive Genetic Variation in Schizophrenia Risk Is Shared by Populations of African and European Descent. Am J Hum Genet 93: 463–470.

Deng Y, Pan W. 2017. Testing Genetic Pleiotropy with GWAS Summary Statistics for Marginal and Conditional Analyses. Genetics 207: 1285–1299.

Duforet-Frebourg N, Luu K, Laval G, Bazin E, Blum MGB. 2016. Detecting Genomic Signatures of Natural Selection with Principal Component Analysis: Application to the 1000 Genomes Data. Mol Biol Evol 33: 1082–1093.

Field Y, Boyle EA, Telis N, Gao Z, Gaulton KJ, Golan D, Yengo L, Rocheleau G, Froguel P, McCarthy MI, et al. 2016. Detection of human adaptation during the past 2000 years. Science.

Fisher RA. 1930. The genetical theory of natural selection. At the Clarendon Press.

Fraser HB. 2013. Gene expression drives local adaptation in humans. Genome Res 23: 1089–1096.

Galinsky KJ, Bhatia G, Loh P-R, Georgiev S, Mukherjee S, Patterson NJ, Price AL. 2016. Fast Principal-Component Analysis Reveals Convergent Evolution of ADH1B in Europe and East Asia. Am J Hum Genet 98: 456–472.

Gratten J, Visscher PM. 2016. Genetic pleiotropy in complex traits and diseases: implications for genomic medicine. Genome Med 8. http://genomemedicine.biomedcentral.com/articles/10.1186/s13073-016-0332-x (Accessed September 22, 2016).

Halko N, Martinsson PG, Tropp JA. 2011. Finding Structure with Randomness: Probabilistic Algorithms for Constructing Approximate Matrix Decompositions. SIAM Rev 53: 217–288.

Han B, Pouget JG, Slowikowski K, Stahl E, Lee CH, Diogo D, Hu X, Park YR, Kim E, Gregersen PK, et al. 2016. A method to decipher pleiotropy by detecting underlying heterogeneity driven by hidden subgroups applied to autoimmune and neuropsychiatric diseases. Nat Genet 48: 803–810.

Hennekens CH, Hennekens AR, Hollar D, Casey DE. 2005. Schizophrenia and increased risks of cardiovascular disease. Am Heart J 150: 1115–1121.

Hermisson J, Pennings PS. 2005. Soft sweeps: molecular population genetics of adaptation from standing genetic variation. Genetics 169: 2335–2352.

Hernandez RD, Kelley JL, Elyashiv E, Melton SC, Auton A, McVean G, 1000 Genomes Project, Sella G, Przeworski M. 2011. Classic selective sweeps were rare in recent human evolution. Science 331: 920–924.

Hine E, McGuigan K, Blows MW. 2014. Evolutionary Constraints in High-Dimensional Trait Sets. Am Nat 184: 119–131.

Johnson T, Barton N. 2005. Theoretical models of selection and mutation on quantitative traits. Philos Trans R Soc Lond B Biol Sci 360: 1411–1425.

Kingsolver JG, Hoekstra HE, Hoekstra JM, Berrigan D, Vignieri SN, Hill CE, Hoang A, Gibert P, Beerli P. 2001. The strength of phenotypic selection in natural populations. Am Nat 157: 245–261.

Klimentidis YC, Abrams M, Wang J, Fernandez JR, Allison DB. 2011. Natural selection at genomic regions associated with obesity and type-2 diabetes: East Asians and sub-Saharan Africans exhibit high levels of differentiation at type-2 diabetes regions. Hum Genet 129: 407–418.

Lande R. 1980. The Genetic Covariance Between Characters Maintained by Pleiotropic Mutations. Genetics 94: 203–215.

Lande R, Arnold SJ. 1983. The Measurement of Selection on Correlated Characters. Evolution 37: 1210–1226.

Majumdar A, Haldar T, Bhattacharya S, Witte J. 2017. An efficient Bayesian meta-analysis approach for studying cross-phenotype genetic associations. http://biorxiv.org/lookup/doi/10.1101/101543 (Accessed June 23, 2017).

McGuigan K, Collet JM, Allen SL, Chenoweth SF, Blows MW. 2014. Pleiotropic Mutations Are Subject to Strong Stabilizing Selection. Genetics 197: 1051–1062.

Messer PW, Petrov DA. 2013. Population genomics of rapid adaptation by soft selective sweeps. Trends Ecol Evol 28. https://www.ncbi.nlm.nih.gov/pmc/articles/PMC3834262/ (Accessed January 24, 2018).

Miura S, Zhang Z, Nei M. 2013. Random fluctuation of selection coefficients and the extent of nucleotide variation in human populations. Proc Natl Acad Sci 110: 10676–10681.

Okbay A, Beauchamp JP, Fontana MA, Lee JJ, Pers TH, Rietveld CA, Turley P, Chen G-B, Emilsson V, Meddens SFW, et al. 2016. Genome-wide association study identifies 74 loci associated with educational attainment. Nature 533: 539–542.

Orchard TR. 2012. Management of Arthritis in Patients with Inflammatory Bowel Disease. Gastroenterol Hepatol 8: 327–329.

Orr HA. 2000. Adaptation and the cost of complexity. Evol Int J Org Evol 54: 13–20.

Patterson N, Price AL, Reich D. 2006. Population Structure and Eigenanalysis. PLoS Genet 2: e190.

Pickrell JK, Berisa T, Liu JZ, Ségurel L, Tung JY, Hinds DA. 2016. Detection and interpretation of shared genetic influences on 42 human traits. Nat Genet 48: 709–717.

Pickrell JK, Coop G, Novembre J, Kudaravalli S, Li JZ, Absher D, Srinivasan BS, Barsh GS, Myers RM, Feldman MW, et al. 2009. Signals of recent positive selection in a worldwide sample of human populations. Genome Res 19: 826–837.

Price AL, Patterson NJ, Plenge RM, Weinblatt ME, Shadick NA, Reich D. 2006. Principal components analysis corrects for stratification in genome-wide association studies. Nat Genet 38: 904–909.

Pritchard JK, Di Rienzo A. 2010. Adaptation - not by sweeps alone. Nat Rev Genet 11: 665–667.

Pritchard JK, Pickrell JK, Coop G. 2010. The genetics of human adaptation: hard sweeps, soft sweeps, and polygenic adaptation. Curr Biol CB 20: R208–215.

Przeworski M, Coop G, Wall JD. 2005. The signature of positive selection on standing genetic variation. Evol Int J Org Evol 59: 2312–2323.

Purcell S, Chang C. PLINK. https://www.cog-genomics.org/plink2.

Robinson MR, Hemani G, Medina-Gomez C, Mezzavilla M, Esko T, Shakhbazov K, Powell JE, Vinkhuyzen A, Berndt SI, Gustafsson S, et al. 2015. Population genetic differentiation of height and body mass index across Europe. Nat Genet 47: 1357–1362.

Rockman MV. 2012. The QTN program and the alleles that matter for evolution: all that’s gold does not glitter. Evol Int J Org Evol 66: 1–17.

Sabeti PC, Reich DE, Higgins JM, Levine HZP, Richter DJ, Schaffner SF, Gabriel SB, Platko JV, Patterson NJ, McDonald GJ, et al. 2002. Detecting recent positive selection in the human genome from haplotype structure. Nature 419: 832–837.

Sabeti PC, Schaffner SF, Fry B, Lohmueller J, Varilly P, Shamovsky O, Palma A, Mikkelsen TS, Altshuler D, Lander ES. 2006. Positive natural selection in the human lineage. Science 312: 1614–1620.

Sanjak JS, Sidorenko J, Robinson MR, Thornton KR, Visscher PM. 2018. Evidence of directional and stabilizing selection in contemporary humans. Proc Natl Acad Sci U S A 115: 151–156.

Schrider DR, Kern AD. 2017. Soft Sweeps Are the Dominant Mode of Adaptation in the Human Genome. Mol Biol Evol 34: 1863–1877.

Simons YB, Bullaughey K, Hudson RR, Sella G. 2018. A population genetic interpretation of GWAS findings for human quantitative traits. PLOS Biol 16: e2002985.

Visscher PM, Yang J. 2016. A plethora of pleiotropy across complex traits. Nat Genet 48: 707.

Vitti JJ, Grossman SR, Sabeti PC. 2013. Detecting Natural Selection in Genomic Data. Annu Rev Genet 47: 97–120.

Walsh B, Blows MW. 2009. Abundant Genetic Variation + Strong Selection = Multivariate Genetic Constraints: A Geometric View of Adaptation. Annu Rev Ecol Evol Syst 40: 41–59.

Wang L, Oehlers SH, Espenschied ST, Rawls JF, Tobin DM, Ko DC. 2015. CPAG: software for leveraging pleiotropy in GWAS to reveal similarity between human traits links plasma fatty acids and intestinal inflammation. Genome Biol 16: 190.

Welch JJ, Waxman D. 2003. Modularity and the cost of complexity. Evol Int J Org Evol 57: 1723–1734.

Wood AR, Esko T, Yang J, Vedantam S, Pers TH, Gustafsson S, Chu AY, Estrada K, Luan J, Kutalik Z, et al. 2014. Defining the role of common variation in the genomic and biological architecture of adult human height. Nat Genet 46: 1173–1186.

Yang J, Benyamin B, McEvoy BP, Gordon S, Henders AK, Nyholt DR, Madden PA, Heath AC, Martin NG, Montgomery GW, et al. 2010. Common SNPs explain a large proportion of the heritability for human height. Nat Genet 42: 565–569.

Yang JJ, Williams LK, Buu A. 2017. Identifying Pleiotropic Genes in Genome-Wide Association Studies for Multivariate Phenotypes with Mixed Measurement Scales. PLOS ONE 12: e0169893.

Zeng J, de Vlaming R, Wu Y, Robinson MR, Lloyd-Jones LR, Yengo L, Yap CX, Xue A, Sidorenko J, McRae AF, et al. 2018. Signatures of negative selection in the genetic architecture of human complex traits. Nat Genet. http://www.nature.com/articles/s41588-018-0101-4 (Accessed April 24, 2018).

